# Measurement of Real Time Serotonin Dynamics from Human Derived Gut Organoids

**DOI:** 10.1101/2024.11.08.622640

**Authors:** B. Bohl, Y. Lei, G A. Bewick, P. Hashemi

## Abstract

The importance of the gut in regulating the brain-body-immune axis is becoming increasingly evident. Interestingly, the brain and gut share many common signalling molecules, with serotonin being one of the most notable. In fact, the gut is the primary source of serotonin in the body. However, studying serotonin dynamics in a human-specific context remains a challenge. Human stem cell-derived models provide a promising avenue for studying signal transmission in well-controlled, *in vitro* environments. In this study, we report the first fast-scan cyclic voltammetry (FSCV) measurements of serotonin signalling in a newly developed enterochromaffin cell (ECC)-enriched gut organoid model. First, we characterize the stem cell-derived gut organoids and confirmed they are enriched with ECCs - the key cell type responsible for producing and releasing serotonin in the gut. We then optimize an *in vitro* buffer that maintains cell viability while supporting FSCV measurements. Using this system, we detect spontaneous release events, which increase in frequency and amplitude following stimulation with forskolin (FSK) and 3-isobutyl-1-methylxanthine (IBMX). Finally, we confirm the identity of the signal as serotonin by using a selective serotonin reuptake inhibitor (SSRI), which significantly delayed the reuptake profile. Our study introduces the first real time measurement of serotonin signalling in a human-derived gut model. We believe this system will be essential for future research on serotonin’s role in the gut and for potential novel drug target identification.

## Introduction

The gut is a critically important area of biological study due to its complex and dynamic functions. Key players in this regulation are the hormone and neurotransmitter producing enteroendocrine cells (ECCs) which are scattered throughout the gut epithelium. These cells act as central communicators, translating luminal, neural, and immune signals into functional responses, locally and body wide. They regulate a broad spectrum of physiological processes, including maintaining metabolic homeostasis, digestion, and gut-brain communication. Disruption of this signalling has been linked to conditions such as irritable bowel syndrome (IBS) (Crowell, 2004; Sikander *et al*., 2009; Vahora *et al*., 2020) and mental health disorders (Coppen, 1967; Cui *et al*., 2024). The gut and brain are connected through a bidirectional immune axis, making gut inflammation and depression well-known comorbidities (Staudacher *et al*., 2023). Interestingly, several neurotransmitters that regulate mood in the brain also exist at high concentrations in endocrine cells in the gut. For instance, enterochromaffin cells are a subset of the enteroendocrine population and produce serotonin (El-Merahbi *et al*., 2015).

In the gastrointestinal tract serotonin regulates peristalsis, motility, fluid and mucus secretion, and nutrient absorption (Hansen and Witte, 2008; Kendig and Grider, 2016; Martin *et al*., 2017). Elevated serotonin levels are observed in the colon and ileum during inflammatory bowel disease (IBD) and in response to bacterial stimulation (Nzakizwanayo *et al*., 2015; Perez *et al*., 2022), whilst disruptions in serotonin signalling have also been implicated in irritable bowel syndrome (IBS) and gastrointestinal motility disorders (Sikander *et al*., 2009). Beyond local effects on the GI, serotonin plays a crucial role in the gut-brain axis where it is thought to modulate mood, stress response, and neuro-gastroenterological disorders (Khlevner *et al*., 2018). Dysregulation of serotonin signalling may contribute to these conditions by altering immune cell responses, the gut microbiome, and epithelial barrier integrity (Chin *et al*.; Fung *et al*., 2017; Keszthelyi *et al*., 2014; Wu *et al*., 2019). Targeting serotonin pathways presents potential novel therapeutic opportunities for many GI disorders. However, there is a critical need to develop more physiologically relevant and specific tools for studying serotonin’s role in the gut.

Over the past decade, electrochemical techniques have offered the first insights into the dynamics of serotonin secretion in the gut. Patel *et al*. (2008) utilized boron doped diamond electrodes to simultaneously measure serotonin and melatonin from the mucosa in the rabbit ileum showing important mechanistic differences between regulation of the two modulators. Amperometric measurements in single human neuroendocrine carcinoid BON cells revealed that 80% of vesicular contents are released during signalling (Wang *et al*., 2021). In both primary cell preparations and cancer cell lines, amperometry demonstrated that although these cells express proteins related to neuronal release machinery, their signal kinetics resemble those of endocrine, rather than neuronal, signalling (Shaaban *et al*., 2021). Yeoman *et al*. (2024) investigated serotonin signalling in the mouse colon and ileum, and discovered regional differences were due to variations in serotonin autoreceptor and serotonin transporter density. Development of flexible electrochemical probes enabled the first *in vivo* amperometric measurements in mice, with minimal impact on tissue integrity or function (Li *et al*., 2022). Recently, fast-scan cyclic voltammetry (FSCV) has been applied to *ex vivo* gut slices for the first time, revealing the simultaneous release of serotonin and melatonin (Delong *et al*., 2024). FSCV can differentiate between various chemical compounds—such as neurotransmitters (*e*.*g*., serotonin, dopamine), hormones, or metabolites— because of cyclic voltammograms (CVs) with discrete oxidation and reduction potentials, afforded by the technique.

Human-derived models offer an opportunity to study human-specific phenomena in well-controlled, *in vitro* environments. The most accessible approach to achieving this is through stem cell-derived models. Despite the significance of ECCs, studying human ECC biology has been challenging due to their sparse distribution and the difficulties associated with isolating and maintaining primary human enterochromaffin cells in culture. Much of our current knowledge stems from murine models, where fluorescent reporters have enhanced our understanding of ECC distribution, differentiation, and function (Billing *et al*., 2019; Song *et al*., 2023). However, interspecies differences underscore the need for a better understanding of human biology. The recent convergence of organoid technology and CRISPR-Cas9 gene editing has transformed the field. By culturing organoids and introducing fluorescent labels *via* CRISPR-Cas9, we can now visualize and study human ECCs in unprecedented detail. However, such models have not been widely probed using electrochemistry.

In this paper, we generated human CHGA-mNeon-expressing colonoids using CRISPR– Cas9-mediated homology-independent transgenesis (CRISPR-HOT). In this context, CHGA (chromogranin A) serves as a marker for enterochromaffin cells (Bellono *et al*., 2017). We began by confirming colonoids contained the necessary biochemical machinery for synthesizing serotonin. Next, we optimized an *in vitro* buffer to ensure compatibility between the organoid culture conditions and FSCV measurements. We subsequently measured spontaneous events and observed CVs corresponding to serotonin, which increased in frequency and amplitude when stimulated with forskolin (FSK) and 3-isobutyl-1-methylxanthine (IBMX). Finally, we verified the identity of the signal using a selective serotonin reuptake inhibitor (SSRI), which significantly slowed the reuptake profile of the signal.

In summary, we present a human gut organoid model which can be used to measure real-time secretion of serotonin from ECC cells using FSCV. We believe this model will be invaluable for future studies on the roles of serotonin in human gut health and disease.

## Methods

### Sample collection and cell culture

Human colon organoids (colonoids) were derived from colonoscopy biopsies of a 50-year-old male patient (unique identifier FG) from Kings College Hospital NHS Foundation Trust. Briefly, the biopsies were rinsed in ice-cold phosphate-buffered saline (PBS; D8537, Sigma-Aldrich), and twice in 10 mM 1,4-dithiothreitol (DTT; 10197777001, Sigma-Aldrich) for 5 minutes at room temperature. To release crypts, tissue was incubated rotating in 8 mM EDTA (15575-038, Invitrogen) in PBS for 1 h at 4 °C, and shaken vigorously in cold PBS. Supernatant was centrifuged for 3 min at 400 rcf and the pellet was washed thrice in cold PBS. Crypts were seeded at a density of 200 crypts per 25 μL of Cultrex Basement Membrane Extract (BME) (3536-005-02, Bio-Techne) in Nunc™ Cell-Culture Treated 48-well plates (150687, Thermo Scientific). The BME was allowed to polymerize for 15 minutes at 37°C before adding 250 μL/well of IntestiCult™ Organoid Growth Medium (06010, STEMCELL), supplemented with 100 units/mL Pen-Strep and 10 μM Y-27632 (Y0503, Sigma-Aldrich). Crypts were maintained in a 37°C incubator with 5% CO2, with media changes every other day.

After organoid development, cells were maintained in human colonoid culture medium and passaged by mechanically dissociating colonoid cultures every 7 days and seed at a 1:6 ratio as previously described (Goldspink *et al*., 2020). Maintenance medium, also IFE medium, contained Advanced DMEM/F-12 (12634, Thermo Fisher), 2 mM GlutaMAX (35050061, Gibco), 10 mM HEPES (15630056, Gibco), 100 units/mL Pen-Strep, 1× B27 supplement (17504044, Gibco), 1× N2 supplement (17502048, Gibco), 0.15nM Wnt Surrogate-Fc Fusion Protein (N001, ImmunoPrecise Antibodies), 10% R-spondin-1 conditioned medium (in house), 1% Noggin-Fc fusion protein conditioned medium (N002, ImmunoPrecise Antibodies), 50 ng/mL recombinant human EGF (AF-100-15, PeproTech), 1.25 mM N-Acetylcysteine (A9165, Sigma-Aldrich), 10 nM Gastrin (G9145, Sigma-Aldrich), 500 nM A83-01 (2939, Bio-techne), 100 ng/mL recombinant human IGF-1 protein (590904, Biolegend) and 50 ng/mL recombinant human FGF-2 protein (100-18B, PeproTech).

Differentiation of colonoids was initiated on day 4 post-passaging by replacing the IFE medium with IF* medium until day 11, reducing the concentration of Wnt Surrogate protein to 0.045 nM, and removing EGF. From day 7, a 48 h treatment with 10 μM Notch inhibitor DAPT (D5942, Sigma-Aldrich), 500 nM MEK inhibitor PD0325901 (Sigma-Aldrich) and 40 μM ISX-9 (4439/10, Bio-Techne) was initiated to boost differentiation. Final maturation was performed in IF* medium and all analysis were performed on day 11.

### Generation of human CHGA-mNeon reporter organoids

CRISPR-HOT technology was used to generate the CHGA-mNeon reporter organoids (Artegiani *et al*., 2020; Beumer *et al*., 2020). Three plasmids– pSPgRNA (Addgene #47108), pCas9-mCherry-Frame +1 (Addgene #66940), and pCRISPaint-mNeon (Addgene #174092) containing an EF1α promoter-driven puromycin resistance gene–were transfected into human colonoids *via* electroporation as previously described (Gaebler *et al*., 2020). Electroporated cells were cultured in IFE medium and selected with 1 μg/mL puromycin after five days.

### Immunofluorescent staining of organoids

The staining of human organoids was performed as previously described (Tsakmaki *et al*., 2020). Briefly, after fixation, blocking and permeabilization, the organoids were incubated with primary antibodies 5-HT (1:200, goat; 20079, Immunostar) overnight at 4°C. On the next day, the organoids were washed and incubated with secondary antibodies (1:500, Alexa Fluor™, Invitrogen) for 1 hr at room temperature. The organoids were mounted with Fluoromount-G® (0100-01, Cambridge Bioscience) and air-dried in the dark.

### Electrode preparation and FSCV measurement

The fabrication of carbon fiber microelectrodes (CFM) and the acquisition of FSCV data were carried out as previously described (Holmes *et al*., 2022). Briefly, a T-650 carbon fiber (7 μm diameter; Goodfellow) was drawn into a glass capillary (1.0 mm outer diameter, 0.58 mm inner diameter; World Precision Instruments) and pulled using a PE-22 micropipette puller (Narishige Group) to form a tight seal. The exposed fiber was manually cut to a length of 100 μm (+/-2 μm) and connected to a pinned stainless-steel wire utilized silver paint. The fiber was coated with NafionTM (Liquion Solution, LQ-1105, 5% by weight Nafion, Ion Power) by applying 1 V (vs. Ag/AgCl) for 30 seconds.

Data collection was conducted using WCCV 3.06 software (Knowmad Technologies), a Dagan potentiostat (Dagan Corporation), and a Pine Research headstage (Pine Research Instrumentation). A waveform optimized for serotonin detection (0.2 V to 1.0 V to -0.1 V to 0.2 V vs. Ag/AgCl at 1000 V/s) was applied. After equilibrating the electrode surface through cycling 10 min at 60 Hz and 10 min at 10 Hz, measurements were taken at 10 Hz with 30 seconds per file.

### Calibrations and stability analysis

Calibrations and stability analysis were performed using a custom-built flow cell as previously described (Hexter *et al*., 2023). Briefly, serotonin solutions of different concentrations were injected into the flow cell for a 10 sec, rectangular analyte pulse. Current response at the CFM was correlated to serotonin concentration and the linear concentration range was fitted.

The following buffer/media compositions were test in the flow cell: Advanced DMEM/F12 (12634, Thermo Fisher) supplemented with 2 mM GlutaMAX (35050061, Gibco), 10 mM HEPES (15630056, Gibco), 100 units/mL Pen-Strep, 1× B27 supplement (17504044, Gibco) and 1× N2 supplement (17502048, Gibco). Secretion buffer containing 138 mM NaCl, 4.5 mM KCl, 4.2 mM NaHCO3, 1.2 mM NaH2PO4, 2.6 mM CaCl2, 1.2 mM MgCl2, and 10 mM HEPES (pH 7.4) with/without 0.1% bovine serum albumin (BSA; 15260037, Gibco).

The stability of serotonin signal over 60 min was test with repeated injections of 100 nM serotonin in secretion buffer without BSA every 2 min.

### Colonoid measurements

Organoids were placed in 3.5cm-dish in 2 ml secretion buffer -BSA on a Bio Station IM (Nikon). GFP-reporter was used to identify ECC-containing organoid structures and CFM was positioned using a QUAD micromanipulator (Sutter Instruments). The set-up was cover with aluminium foil connected to the ground as a Faraday cage. After cycling, 15 min of back-to-back files were taken, before adding 10 μM Forskolin (FSK) and 10 μM 3-isobutyl-1-methylxanthine (IBMX) and recording another 15 min. For SSRI treatment, 10 uM Escitalopram (Sigma Aldrich) was added after the initial 15 min, incubated for 5 min, before adding FSK/IBMX.

### Analysis

Data analysis and export of FSCV data was performed using WCCV 3.06 software (Knowmad Technologies). Graphs were generated and statistical analysis were performed using Prism10 (GraphPad). All quantifications are shown as mean +/-standard error of the mean (SEM), unless otherwise stated. Significance levels were set as follows: * P < 0.05, ** P < 0.005, and *** P < 0.001. Number of replicates and statistical tests are indicated in each figure legend. Schematic representations were generated using biorender.com.

## Results and Discussion

### An Optimized Buffer to Facilitate Cell Functionality and FSCV Measurements

When optimizing electro-analytical measurements in novel biological systems, it is important to characterize the electrochemical response in the cell media specific for the cell type being studied. This is because these media contain a complex variety of substrates and proteins that could interact negatively with the sensors. For example, in previous work we found that we could not use a common culture medium for neurons, ‘Neurobasal Medium’, to measure serotonin from human derived serotonergic neurons with FSCV (Holmes *et al*., 2022). Exposure of this medium to the electrode dramatically decreased the analytical response to serotonin. In this previous work, we found that we could keep the cells alive for the duration of the experiment in HEPES buffer than included glucose (Holmes *et al*., 2022). Here, we performed a similar experiment where we utilized advanced Dulbecco’s Modified Eagle Medium (advanced DMEM), which is the colonoid culture medium, as the flow injection buffer during an FSCV injection of serotonin (500 nM). **Figure 1A** shows the FSCV colour plot of this injection. Interpretation of these colour plots can be aided *via* reference to Michael *et al*. (1999). Briefly, voltage is plotted on the y axis with time on the x axis and current in false colour. The absence of a stereotypical CV response to serotonin shows that this medium, as with the ‘Neurobasal Medium’ has reduced the capacity of the electrode to respond to serotonin. This is likely because of fouling of the surface by amino acids in the medium that have the capacity to electropolymerize on the electrode surface (Holmes *et al*., 2022; Jang *et al*., 2024; Shin *et al*., 2005; Takmakov *et al*., 2010). We quantified the peak at approximately 0.6 V (where we expect to capture serotonin oxidation) and found 0.92 +/-0.27 nA. When we repeated the experiment in ‘Secretion Buffer’ used routinely for secretion analysis of gut organoids by mass spectroscopy (Miedzybrodzka *et al*., 2020), we captured a typical serotonin CV, and a much higher signal of 7.63 +/-0.23 nA (**Figure 1B**).

**Figure 1:**
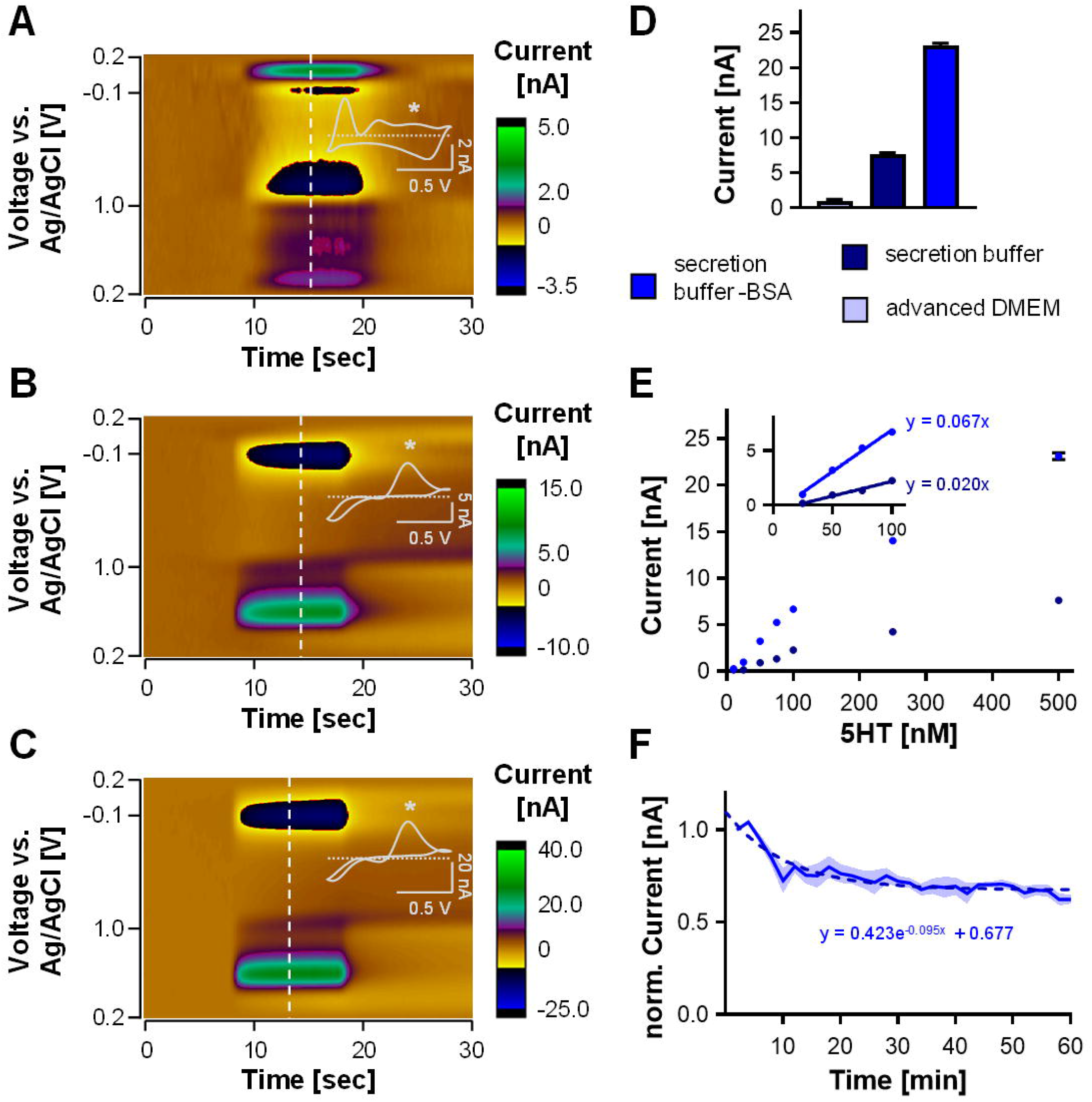
FSCV buffer optimization. **(A-C)** Representative color plot for 500 nM serotonin injections in advanced DMEM (A), secretion buffer (B) and secretion buffer -BSA (C) with CV inset showing serotonin specific oxidation peak (*). **(D)** Peak current measured from 500 nM serotonin injections in the different buffers/media (n = 9 measurements on 3 electrodes, mean +/-SEM). **(E)** Calibration curve for secretion buffer with and without BSA showing serotonin concentration vs. peak current and regression of the linear range in the inset (n = 3 for each buffer, mean +/-SEM). **(F)** Stability analysis for 100 nM serotonin injections in secretion buffer -BSA over a total of 60 min with exponential decay fitting as dashed line (n = 4, mean +/-SEM).

Secretion buffer contains 0.1% bovine albumin serum (BSA) which prevents attachment of organoids and analytes to plastic surfaces. We find that presence of BSA reduces the FSCV serotonin signal, again, likely because of fouling of the electrode by BSA’s electroactive amino acid residues. Auspiciously, for this type of real-time, end-point FSCV experiments, this attachment is not an important consideration thus BSA can be excluded from the buffer. When removing the BSA, the serotonin signal was 23.11 +/-0.39 nA (**Figure 1C**), which is comparable to signals we have seen before in TRIS buffer (Hexter *et al*., 2023). **Figure 1D** shows the pooled responses of the three buffers and **Figure 1E** compares calibration curves between the ‘secretion buffer’ with and without BSA. Calibrations show that without the BSA, the response of the electrode is more sensitive with slopes of linear regression of 0.076 +/-0.003 nA nM^-1^ (secretion buffer -BSA) and 0.027 +/-0.001 nA nM^-1^ (secretion buffer).

Given the superior response in BSA free buffer, we assessed the stability of the response to repeated injections of serotonin and found that the electrode reached stability in a profile that could be fit with an exponential decay over 60 minutes (**Figure 1E**), which is consistent with stability of the serotonin signal in TRIS buffer (Hexter *et al*., 2023). Therefore, having established the suitability of the ‘Secretion Buffer minus BSA’ for FSCV experiment, we proceeded with our physiological experiments using this buffer.

### Characterization of the Colonoids Reveals the Presence of Serotonin-producing ECCs

Previous work using fluorescent reporters has greatly contributed to our knowledge of ECC distribution, differentiation, and function but is derived from murine models (Billing *et al*., 2019; Shajib *et al*., 2013; Song *et al*., 2023). Therefore, as an *in vitro*-model system for the human gut, we derived human colon organoids (colonoids) from primary colonic crypts. The expansion and differentiation protocol, designed to foster ECC development (**Figure 2A**) is described in detail elsewhere (Lei *et al*., 2024). To identify endocrine structures, we used the expression of our recently developed mNeonGreen reporter under the control of the Chromogranin A (CHGA) promoter. Immunostainings confirmed the enrichment of ECCs synthesizing serotonin in mNeonGreen-positive structures (**Figure 2B**), while a smaller fraction of mNeonGreen-positive cells express glucagon-like peptide 1 (GLP-1) as a marker for enteroendocrine cells (**Figure 2C**). Therefore, the mNeonGreen-reporter represents a valuable tool to identify serotonin-producing ECCs in living colonoids.

**Figure 2:**
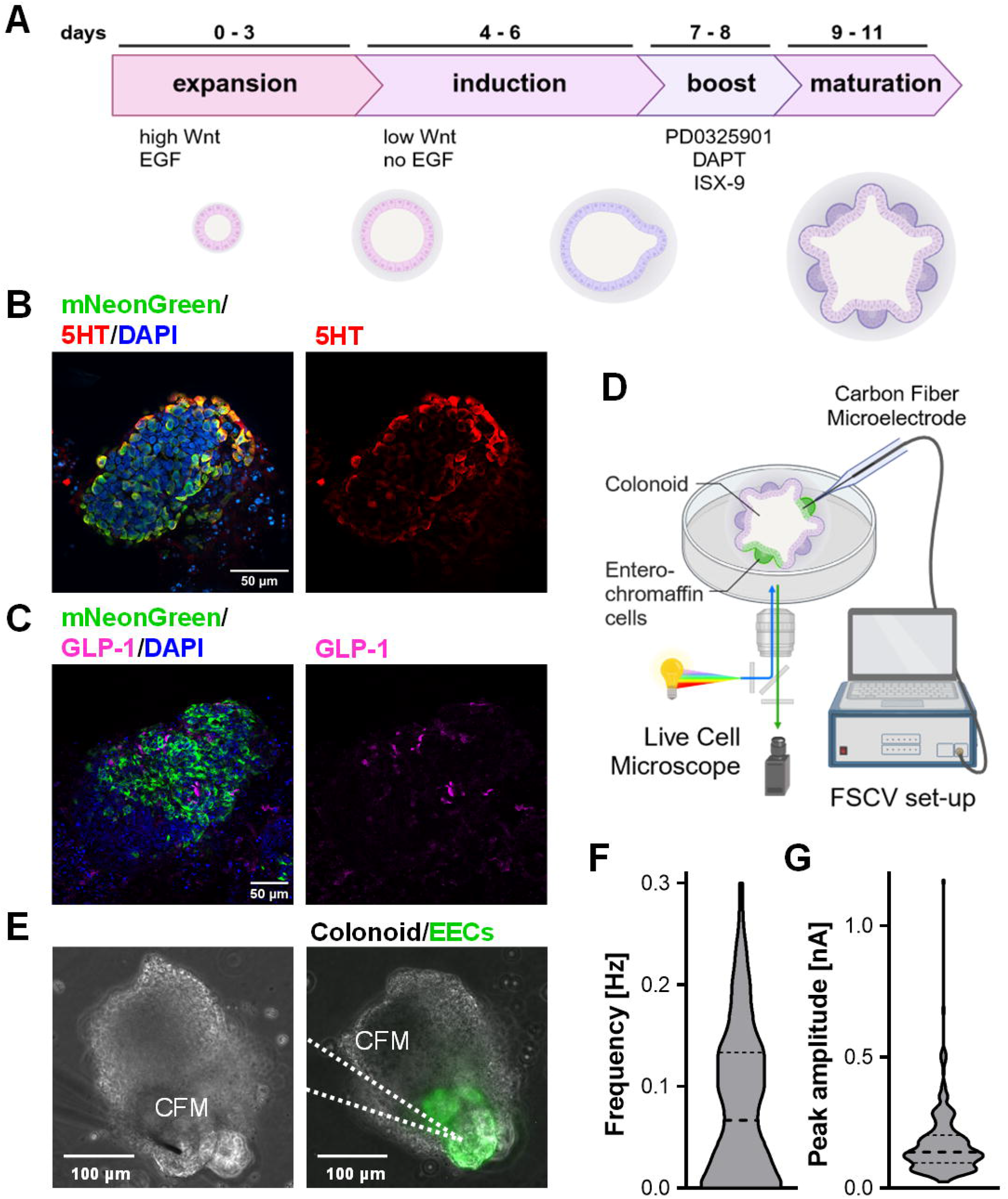
Characterization of serotonin secreting cells in human colonoids by immunofluorescence and FSCV. **(A)** Schematic showing the differentiation protocol of human colonoids for the enrichment of enteroendocrine cells (EECs). **(B-C)** Whole-mount immunostaining of fixed CHGA-mNeonGreen human colonoids, stained for ECC synthesizing 5-HT (B) and enteroendocrine marker GLP-1 (C) with Hoechst nuclear counterstain (blue). Scale bar: 50 μm. **(D)** Scheme of FSCV set-up for colonoid measurements. **(E)** Dish containing colonoids are placed on Live Cell microscope to identify mNeonGreen-expressing structures and provide temperature control. Scale bar: 100 μm. **(F)** Frequency of spontaneous serotonin release events (n = 90 files from 3 organoids, violin plot with median and quantiles as dash lines). **(G)** Peak amplitude of serotonin releases (n = 220 events from 3 organoids, violin plot with median and quantiles as dash lines).

Chemical measurements from human derived models are an emerging research tool, providing invaluable insights in the functionality of the derived somatic cells. Measurement of neurotransmitter production from iPSC-derived neurons has been reported for dopaminergic (Zanetti *et al*., 2021) as well as serotonergic neurons (Holmes *et al*., 2022; Nakatsuka *et al*., 2023). A first study employing differential pulse voltammetry was used to study variability in the development of human kidney organoids (Suhito *et al*., 2022). We immersed the colonoids in BSA free ‘Secretion Buffer’ in a dish on a LiveCell microscope (**Figure 2D**). We identified the ECCs *via* the expression of the mNeonGreen reporter and placed the electrode on the surface of the ECC-containing structures (**Figure 2E**). After 20 minutes of equilibration time, we took FSCV files for 15 minutes. We observed sparse but robust spontaneous electrochemical events during this control time with a mean frequency of 0.082 +/-0.00 Hz and a mean amplitude of 0.169 +/-0.00 nA (**Figure 2F-G**). The frequency of these events mirrors those seen previously in stimulated BON cells (Wang *et al*., 2021).

We provide here evidence of spontaneous secretion from the human derived organoids. The CVs of these events strongly resemble FSCV serotonin (Holmes *et al*., 2022; Wood *et al*., 2014), therefore we next performed some pharmacology to validate the identity.

### Spontaneous Release Events Increase their Frequency and Amplitude with cAMP Agonism

A variety of stimuli have been employed to release serotonin from previous gut models, including ionomycin, a Ca^2+^ ionophore (Shaaban *et al*., 2021), high extracellular Ca^2+^ (Wang *et al*., 2021) and mechanical (Bhavik and Patel, 2008). In this work, we stimulated these colonoids with an adenylate cyclase activator, FSK, and a phosphodiesterase inhibitor, IBMX, both of which increase the intracellular cyclic adenosine monophosphate (cAMP). cAMP increases the secretory capacity from rat biliary epithelium (Ammon and Müller, 1984) and FSK potentiates insulin secretion from rat islets (Francis *et al*., 2004). The results of this stimulation are shown in **Figure 3**. Representative colour plots and IT curves are shown in **Figure 3A-B** left panel with a signature serotonin CV. These agents increase the intracellular levels of the second messenger cyclic AMP and activate protein-kinase, and in turn are thought to induce serotonin release by exocytosis. The combination of those two agents has been previously described to induce GLP-1 release from enteroendocrine cells *via* intracellular Ca^2+^ signalling (Simpson *et al*., 2020). In line with this, we found a clear increase in the total number of events (by 399.1% +/-394.8%), however with quite some organoid-to-organoid variation (n = 3 organoids, **Figure 3C**). Similarly, the mean number of release events per file increased for each organoid, with significant increases in small and big events below and above 12.5 nM of serotonin released in 2 organoids each, and an increase in prolonged releases longer than 3 sec in one of the organoids (**Figure 3D**). We believe those variations are dependent on the position of the electrode with respect to the density of ECCs within the colonoid structure and the size of the structure. This experiment has demonstrated that colonoids increase their secretion activity when stimulated with FSK/IBMX.

**Figure 3:**
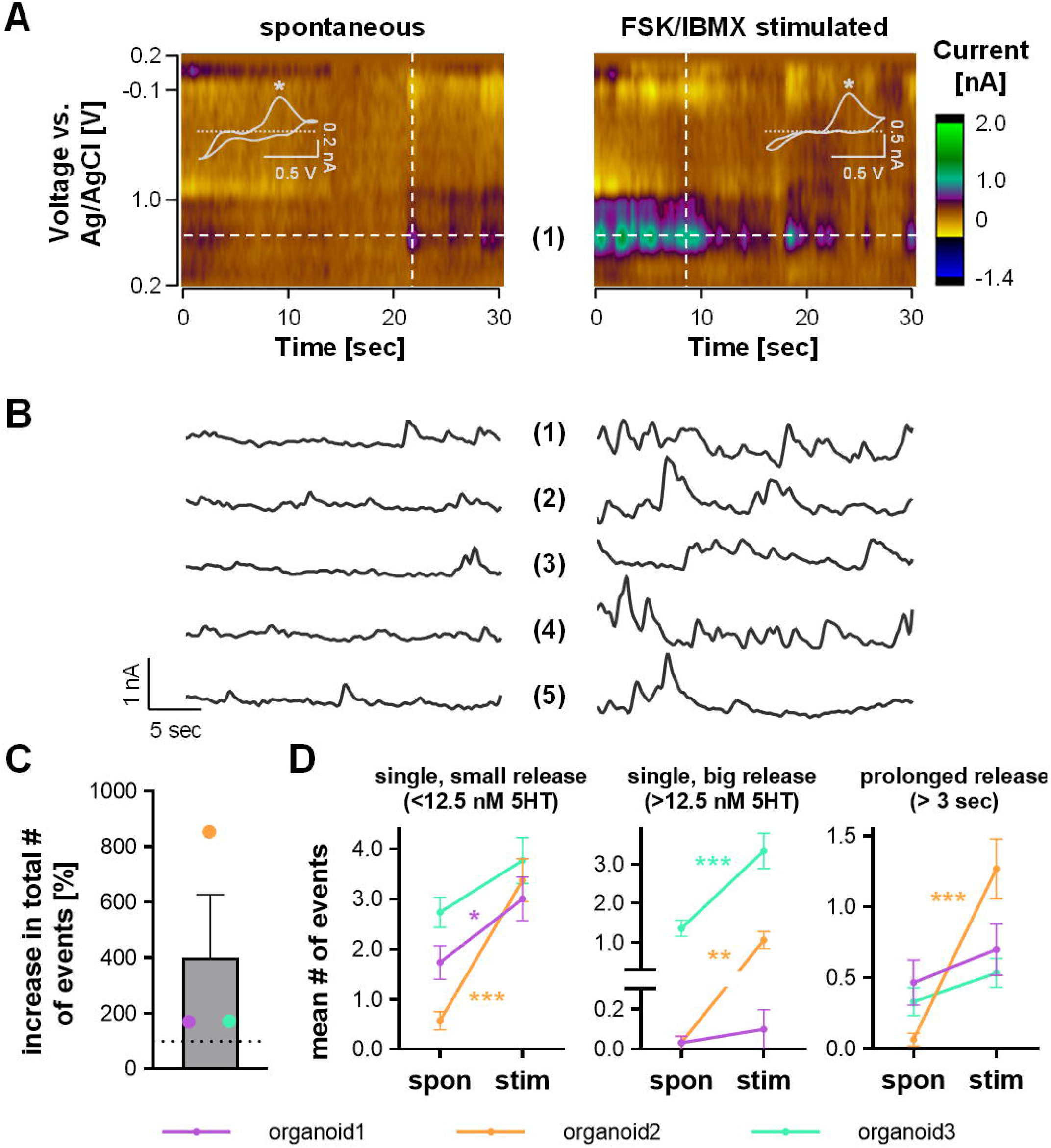
FSCV analysis of spontaneous and stimulated serotonin release from colonoids. **(A)** Representative color plots with CV inset showing serotonin specific oxidation peak (*) and **(B)** IT curves of spontaneous and FSK/IBMX-stimulated releases (10 μM FSK and 10 μM IBMX). **(C)** Percent increase of total number of serotonin release events over 15 min after stimulation (n = 3 organoids, mean +/-SEM). **(D)** Mean number of small (<12.5 nM serotonin), big (>12.5 nM serotonin) and prolonged (>3 sec) release events per file (n = 30 files per organoid; mean +/-SEM; two-way ANOVA with Sidak’s post-hoc test, * p < 0.05, ** p < 0.01, *** p < 0.001).

### Real time Serotonin Signalling from Colonoids is Modulated by Serotonin Transporter Activity

To further validate the chemical identity of the signal, we employed a SERT inhibiting agent. SERTs are transmembrane transporters that reuptake serotonin with high capacity and are inhibited by the popular antidepressants, the SSRIs (Blackburn *et al*., 1967; Coleman *et al*., 2016). We and others have found in the brain that SERT inhibition with SSRI exerts profound effects on the FSCV serotonin signal (Dunham and Venton, 2022; Mena et al., 2024; Witt et al., 2023; Wood and Hashemi, 2013). SERTs are only found in the brain, but are also highly concentrated in the gut (Gill *et al*., 2008). Indeed, SSRIs have clinical utility for treating irritable bowel syndrome (IBS) (Fritsch *et al*., 2020). Therefore, we employed escitalopram (escit) to test the effect on the FSCV signal in the organoids. **Figure 4A** is a colour plot, representative of an isolated FSK/IBMX induced release and **Figure 4B** is a representative plot of a similar event 5 minutes after treatment with escit. Visually, the duration of the signal appears longer after SSRI and to verify this notion, we employed exponential fitting of the reuptake curves across 35 and 19 events for control and escitalopram, respectively (**Figure 4C**). We found a significant decrease in the initial slope (FSK/IBMX = 1.80 ± 0.20 sec^-1^, FSK/IBMX + escit = 1.19 ± 0.13 sec^-1^ indicating the inhibition of reuptake by SERTs (**Figure 4C**,**D**). Interestingly, there was a significant increase in the mean steady state serotonin concentration post event after escit from -0.5732 to 0.1605 ([5HT^0^/5HT^max^]) (**Figure 4C**,**E**). We have previously observed this effect before *in vivo* in the mouse brain which we attributed to SSRI-induced SERT internalization (Lau *et al*., 2008; Witt *et al*., 2023). Based in this finding, an interesting future area of study would be to investigate whether SERT internalization applies to gut epithelial cells.

**Figure 4:**
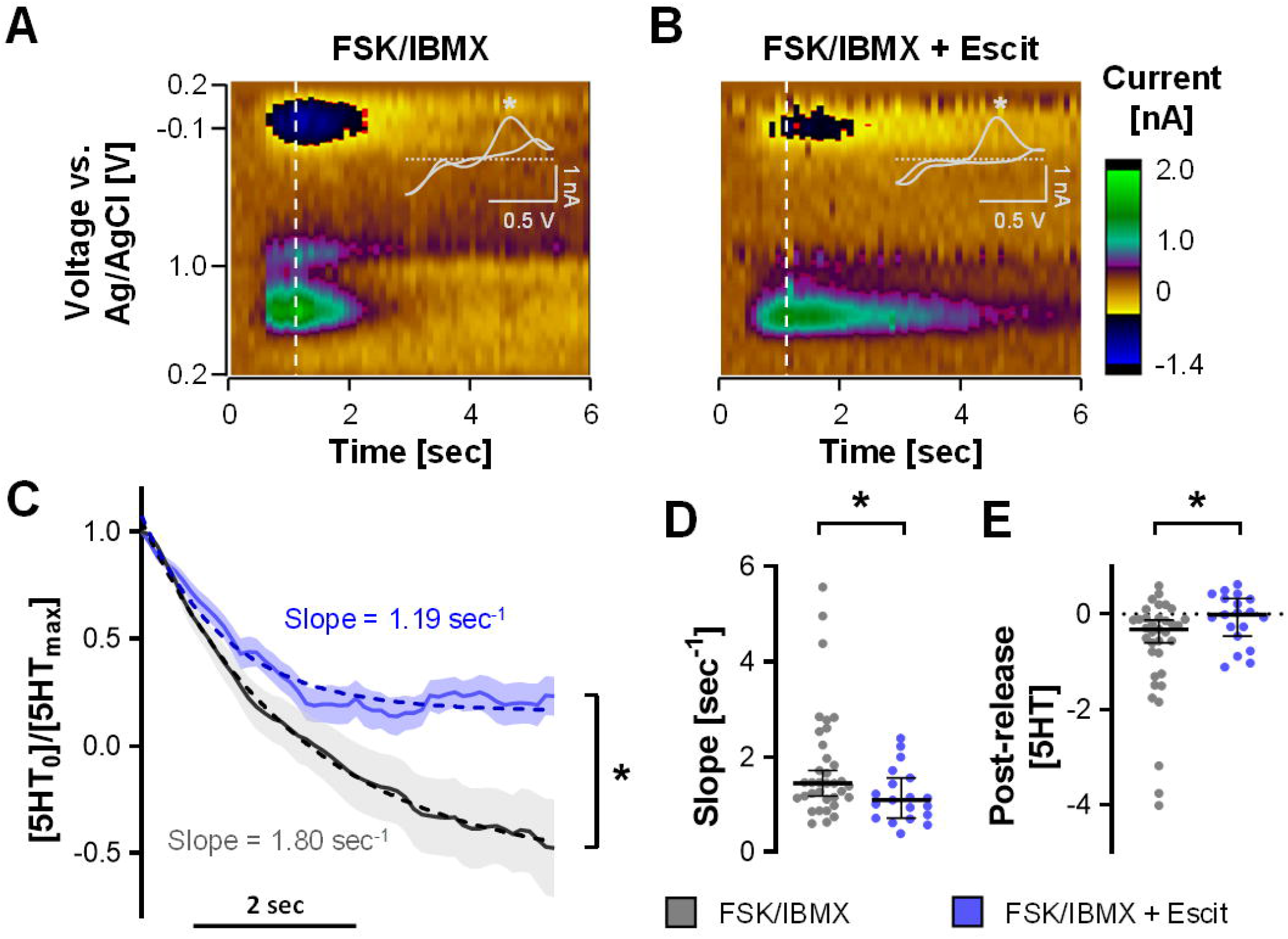
Escitalopram treatment blocks Serotonin reuptake in gut endothelium. **(A-B)** Representative color plots with CV inset showing serotonin specific oxidation peak (*) under FSK/IBMX stimulation condition without (A) and with pre-treatment with 10 uM Escitalopram (Escit) (B). **(C)** Normalized serotonin concentration starting from the peak of release events with (blue) and without (gray) pre-treatment with Escitalopram and exponential decay fitting as dashed lines (FSK/IBMX: n = 35; FSK/IMBX+Escit = 19 events from 3 organoids, mean +/-SEM; two-way ANOVA with Sidak’s post hoc test, * p < 0.05). **(D-E)** Slopes and post-release serotonin concentration calculated from exponential decay fitting of individual release events (FSK/IBMX: n = 33; FSK/IMBX+Escit = 19 events from 3 organoids, median with 95% CI; unpaired t-test, * p < 0.05).

Our study resembles previous work by Yeoman *et al*. who found that SSRIs decreased the slope of their amperometric signal in colonic tissues. As above, SSRIs are clinically prescribed to treat IBS (Fritsch *et al*., 2020). The clinical use of antidepressants for IBS is based on 3 theories (Talley, 2003). The first is that because IBS and depression are highly comorbid (Staudacher *et al*., 2023), the therapeutic effects of SSRIs on depression might in turn reduce the severity of IBS. The second is based on the central analgesic actions of antidepressants and their effects in restoring serotonin homeostasis (Kułak-Bejda *et al*., 2017; Lynch, 2001). Finally, it has previously been postulated that these agents may interact locally with the gut and the intestinal barrier (Ba *et al*., 2024; Eyzaguirre-Velásquez *et al*., 2020; Teng *et al*., 2024). While we cannot localize the precise mechanism with our study of the three theories, we provide compelling evidence for the direct action of SSRIs on gut serotonin.

## Conclusion

The gut is increasingly being recognized to play a central role in modulating the brain-body immune axis. The gut and the brain share many signalling molecules, in particular serotonin. Stem cell derived, human models open interesting avenues for studying human specific phenomena in well controlled, *in vitro* environments, however models such as these have not widely been probed with electrochemistry. Here we described FSCV measurements of serotonin signalling in a newly generated CHGA reporter organoids, where primary human ECCs are fluorescently labelled for easy identification.

We first biochemically characterized the gut organoids and found that they contained the essential cell type producing and releasing serotonin in the gut, ECCs. Then we optimized an *in vitro* buffer for cell culture and FSCV measurements and measured spontaneous events that increased in frequency and amplitude upon stimulation with FSK/IMBX. Finally, the identity of the signal was validated to be serotonin with an SSRI which significantly slowed the reuptake profile of the signal. Therefore, we presented here the first FSCV measurements in an organoid model, and also a new model for measuring serotonin transmission in a human derived gut system that we believe will be invaluable for future studies of the roles of serotonin in gut health and disease.

## Author Contributions

B.B. designed and conducted the majority of experiments, analyzed the data and generated the figures. Y. L. generated the colonoids and carry out molecular characterization. P.H., B.B. and G.B. wrote the manuscript. G.B. and P.H. conceptualized and mentored the work.

## Acknowledgments

This work was supported by Imperial College London and Community for Electroanalytical Measurements. B.B. was supported by the German Research Foundation (BO6292/1-1). Y.L. acknowledges support from China Scholarship Council (K-CSC) and the Society for Endocrinology (Early Career Grant).

